# Growth hormone induces mitotic catastrophe of podocytes and contributes to proteinuria

**DOI:** 10.1101/597344

**Authors:** Rajkishor Nishad, Dhanunjay Mukhi, Ashish Kumar Singh, Kumaraswami Chintala, Prasad Tammineni, Anil Kumar Pasupulati

## Abstract

Podocytes are integral members of the filtration barrier in the kidney and are crucial for glomerular permselectivity. Podocytes are highly differentiated and vulnerable to an array of noxious stimuli during various clinical conditions whereas podocyte loss plays a key role in progressive glomerular diseases. Elevated circulating growth hormone (GH) levels are associated with podocyte injury and proteinuria in diabetics. Previous studies have shown that podocytes express GH receptors (GHR), and induce Notch signaling when exposed to GH. However, the precise mechanism(s) by which excess GH elicits podocytopathy remains to be elucidated. In the present study, we demonstrate that GH induces cognate TGF-β1 signaling and provokes cell cycle re-entry of otherwise quiescent podocytes. Though, differentiated podocytes re-enter the cell cycle in response to GH and TGF-β1 unable to accomplish cytokinesis, despite nuclear division. Owing to this aberrant cell-cycle events significant amount of GH or TGF-β1 treated cells remain binucleated and undergo mitotic catastrophe. Importantly, inhibition of GHR, TGFBR1, or Notch signaling prevented cell cycle re-entry and protects podocyte from cell death. Furthermore, inhibition of Notch activation prevents GH-dependent podocyte injury and proteinuria. Kidney biopsy sections from patients with diabetic nephropathy show activation of Notch signaling and bi-nucleated podocytes. All these data confirm that excess GH induces Notch1 signaling via TGF-β1 and contributes to the mitotic catastrophe of podocytes. This study highlights the role of aberrant GH signaling in the podocytopathy and the potential application of inhibitors of TGF-β1 or Notch inhibitors as a therapeutic agent for diabetic nephropathy.

**Significance Statement:** Elevated circulating levels of growth hormone (GH) associated with glomerular hypertrophy and proteinuria. Whereas decreased GH action protected against proteinuria. Podocytes are highly differentiated cells that play a vital role in glomerular filtration and curb protein loss. The direct role of GH in podocytes is the focus of our study. We found that GH induces TGF-β1 and both provoke cell cycle re-entry of podocytes in Notch1 dependent manner. Notch activation enables the podocytes to accomplish karyokinesis, but not cytokinesis owing to which podocytes remain binucleated. Binucleated podocytes that were observed during GH/TGF-β1 treatment are susceptible to cell death. Our study highlighted the fact that enforcing the differentiated podocytes to re-enter the cell cycle results in mitotic catastrophe and permanent loss.

## Introduction

Glomerular complications are the predominant cause of end-stage kidney disease and clinical conditions such as diabetes and hypertension are associated with glomerular dysfunction and proteinuria. Glomerular podocytes are highly differentiated specialized visceral cells that account for about 30% of glomerular cells. These cells provide epithelial coverage to the capillaries and together with glomerular basement membrane (GBM) and perforated endothelial cells constitute a glomerular filtration barrier (GFB). The unique cytoplasmic extensions of podocytes are known as foot-processes, which attach to the GBM and interdigitate with neighboring foot-processes to form the slit-diaphragm (SD). The sophisticated architecture of SD contributes to glomerular permselectivity. The process of progressive podocyte damage characterized by podocyte hypertrophy, detachment of podocytes, and, finally, irreversible loss of podocytes has been observed in human and experimental models of nephropathy and glomerular diseases [1]. Injury and depletion of podocytes, leading to podocyte insufficiency and capillary collapse, have been implicated in the development of glomerulosclerosis and chronic kidney disease. Since matured podocytes are terminally differentiated and quiescent [2], injured podocytes are hardly replaced and leave denuded areas on the glomerular capillary which results in albuminuria [3, 4].

Albuminuria is a marker for renal dysfunction in the general population and is an early marker for overt nephropathy in diabetic subjects. Elevated circulating growth hormone (GH) levels and increased renal expression of the GH receptor (GHR) are associated with nephropathy in poorly controlled type1 diabetes [5, 6]. Excess GH conditions are characterized by glomerular hypertrophy, sclerosis, and albuminuria, whereas blunting GH action is protected from glomerulopathy [7]. Gaddamedi et al showed that podocytes express GHR and canonical JAK-STAT signaling is activated when podocytes are exposed to GH [8]. Our previous work showed that GH activates Notch signaling [9] and promotes epithelial to mesenchymal transition (EMT) of podocytes by inducing ZEB2 (Zinc Finger E-Box Binding Homeobox 2), also known as smad-interacting protein1 (SIP1) [10, 11].

Interestingly, GH-induced glomerulosclerosis and interstitial fibrosis in diabetic rats is associated with increased TGF-β1 levels [12], whereas inhibition of JAK2, an immediate downstream target of GH reduced TGF-β1 expression [13]. Although multiple studies revealed TGF-β1’s role in morphologic manifestations and clinical characteristics of DN, the stimuli that activate the TGF-β/SMAD pathway in the podocytes remain unclear [13, 14]. In the present study, we demonstrate that GH induces TGF-β1 expression, which in-turn transduces Notch activation. GH and TGF-β1 dependent Notch activation stimulated podocyte re-entry into the cell cycle. Nevertheless, persistent activation of Notch signaling resulted in cytokinesis failure and podocyte apoptosis.

## Results

### 1. GH induces TGF-β1 and cognate TGF-β-SMAD pathway in podocytes

Considering the established role of GH and TGF-β1 in eliciting podocyte injury, we investigated the direct action of GH on the TGF-β/SMAD pathway. TGF-β1 mRNA (Fig.1A&B) and protein (Fig.1C&D) levels were up-regulated in both dose (0-500 ng/ml) and time-dependent (0-48 h) manner in GH treated human podocytes (HPC). Immunofluorescence analysis also revealed GH induced TGFβ1 expression in podocytes (Fig.1E). We also observed GH dependent expression of pSTAT3 (Tyr705) and components of TGF-β1 signaling such as TGFBR1 (TGF-β Receptor 1) and pSMAD2/3 in podocytes (Fig.1C&D) and HEPG2 cells (*SI Appendix*, Fig. S1A&B). Re-analysis of our microarray data (#GEO-GSE21327) revealed the up-regulation of TGF-β1 and its receptor in GH-treated podocytes (*SI Appendix*, Fig.S1C). As TGF-β1 is a secretory molecule, we estimated it levels in conditioned medium from GH (GH-CM) treated podocytes and observed that TGF-β1 is induced by GH in both dose and time-dependent manner (Fig.1F&G). We verified the TGF-β1 activation in GH treated podocytes by performing SMAD4-Luciferase activity assay (Fig.1H&I). Furthermore, podocytes isolated from mice (MPC) administered with GH showed elevated expression of TGFBR1 and its ligand TGF-β1 on mRNA and protein levels (Fig.1J&K) and their enhanced staining in the glomerulus (Fig.1L). TGF-β1 was also detected in urine from mice administered with GH (Fig.1M). To demonstrate the paracrine action of GH induced TGF-β1, we exposed podocytes & HepG2 to GH-CM from respective cell types. Interestingly, GH-CM induced phosphorylation of SMAD2&3 in cells naïve to GH treatment (*SI Appendix*, Fig.S1D&E). We also observed the accumulation of SMAD4 in the nucleus and its enhanced promoter activity following treatment with GH or GH-CM (*SI Appendix*, Fig.S2A&B). Furthermore, the paracrine activity of TGF-β1 in GH-CM was demonstrated using a GFP expression plasmid, which is under the control of SMAD binding element. GH-CM induced GFP expression in this model whereas GH-CM that were pre-incubated with anti-TGF-β1 antibody failed to elicit GFP expression (*SI Appendix*, Fig.S2C). Together, these data support the activation of cognate TGF-β1/SMAD signaling in podocytes by GH.

**Figure 1.**
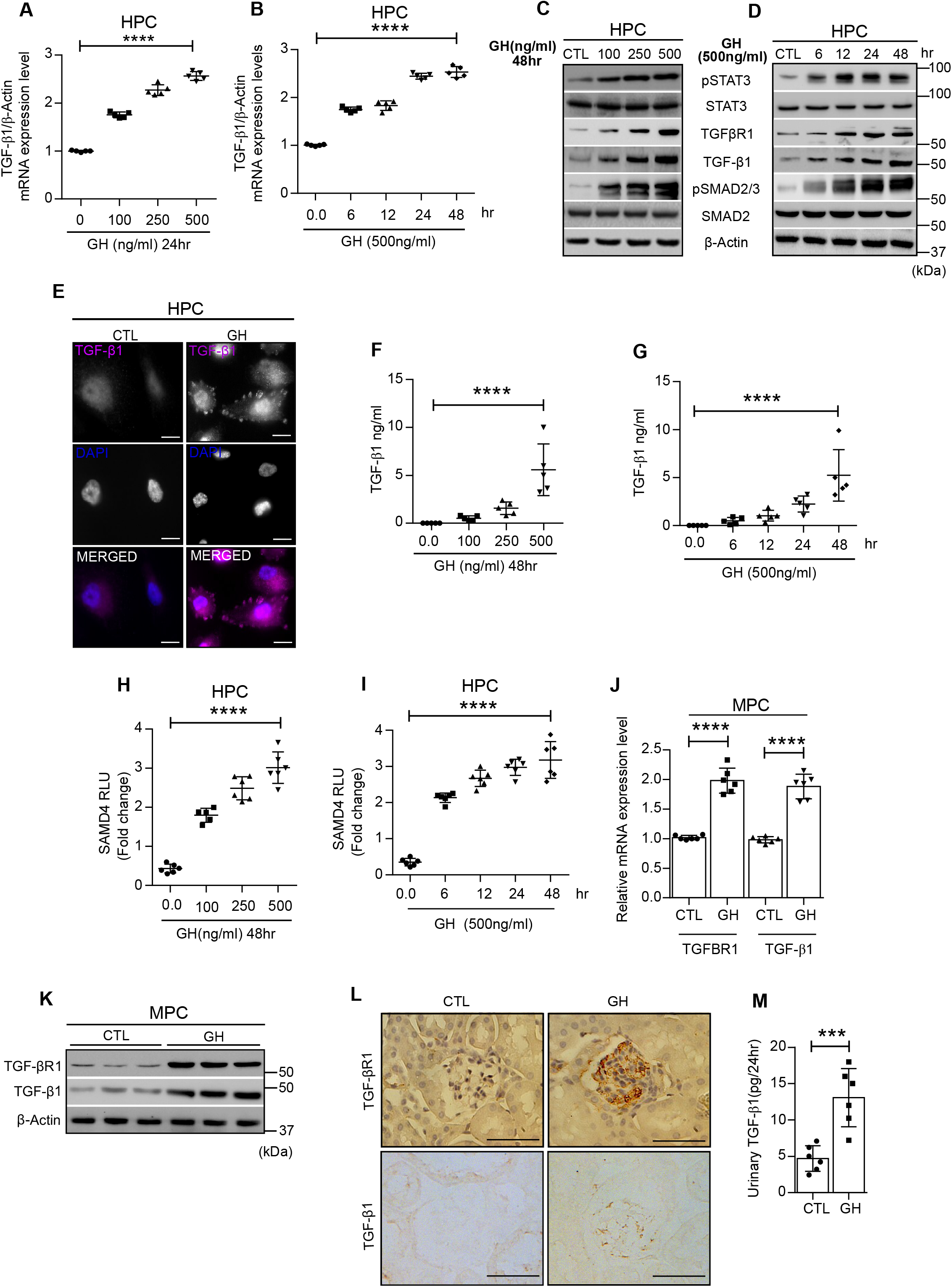
GH induces TGF-β/SMAD pathway in podocytes. **(A&B)** qRT-PCR analysis showing the expression of TGF-β1 in human podocytes (HPC) treated with or without (CTL; control) GH in concentration (100 to 500 ng/ml) and time (up to 48 hr) dependent manner. mRNA levels were normalized to β-Actin and presented as fold-change on y-axis. Mean±SD. (n=5). ****p<0.0001 by 1-way ANOVA post hoc Dunnett test. **(C&D)** Immunoblotting analysis from HPC treated with GH in concentration (100 to 500 ng/ml) and time (up to 48hr) dependent manner. (n=3). **(E)** Immunofluorescence analysis for TGF-β1(purple color) and counter-stained with DAPI [4’,6-diamidino-2-phenylindole (Blue color)] in HPC treated with or without GH (500 ng/ml, 48hr). Scale bar: 50 μm, magnification x500. **(F&G)** Estimation of TGF-β1 in conditioned media (CM) from HPC treated with GH in concentration (100 to 500 ng/ml) and time (up to 48hr) dependent manner. Mean±SD. (n=5). ****p<0.0001 by one-way ANOVA post hoc Dunnet test. **(H&I)** SMAD4 luciferase activity in HPC was treated with GH in concentration (100 to 500 ng/ml) and time (up to 48 hr) dependent manner. Mean±SD. (n=6). ****p<0.0001 by one-way ANOVA post hoc Dunnet test. **(J&K)** qRT-PCR and immunoblotting analysis showing the expression of TGFBR1 and TGF-β1 in CTL or GH-treated (1.5mg/Kg bw) mice podocytes (MPC). Mean±SD. (n=3). ****p<0.0001 by Student t-test. Each dot represents the average value of a single mice from each group (n=6). **(L)** Representative images for TGFBR1 and TGF-β1 expression by DAB staining in mice glomerular sections from CTL vs GH treated mice. Scale bar:100μm, magnification x200. (n=3). **(M)** Quantification of TGF-β1 in urine from CTL or GH treated mice. Mean±SD. (n=3) ****p<0.0001 by Student t-test. Each dot represents the average value of a single mice from each group. β-Actin served as internal control.

### 2. TGF-β1 signaling is required for GH induced Notch reactivation in podocytes

Notch activation was virtually undetectable in glomeruli from heathy adult kidney, unlike their progenitors in the fetal kidney which show enhanced Notch activity [15]. Previously, we showed that GH activates Notch signaling in adult podocytes [9]. Earlier to us, Niranjan et al. 2008 reported that TGF-β1 induces Notch signaling in podocytes from diabetic mice [16]. Since circulating levels of GH elevate in type1 diabetes milieu, and both GH and TGF-β1 were shown to induce Notch signaling we sought to investigate whether GH activates Notch signaling via TGF-β1. When podocytes naive to GH were exposed to GH-CM, activation of Notch signaling was observed similar to that of podocytes treated with GH (Fig.2A&B). It is noteworthy that a TGFBR1 inhibitor (SB431542) ameliorated the effect of GH-CM to induce Notch activation (*SI Appendix*, Fig.S2D). Interestingly, expression of Notch1 and its downstream targets were ameliorated when podocytes were treated with GH in the presence of inhibitors for either GHR (AG490) or TGFBR1 (Fig.2C). Also, GH fails to induce Notch signaling components in engineered podocytes with a specific knockdown of TGFBR1(Fig.2D). Further, the transcriptional activity of NICD1 (Notch intracellular domain) as determined by both nuclear localization of NICD1 and HES1-Luciferase reporter assay is elevated upon GH or TGF-β1 treatment (Fig.2E&F). Nuclear localization of NICD in podocytes primed with GH or TGF-β1 was reduced in the presence of AG490 or SB431542 (Fig.2E). Similarly, the observed increase (~45%) in the transcriptional activity of HES1 by GH or TGF-β1 was abrogated by AG490 or SB431542 (Fig.2F). γ-Secretase is an intracellular protease that cleaves NICD from the Notch receptor and triggers the Notch cascade. Increased γ-secretase activity by GH and TGF-β1 was abolished when podocytes treated with AG490 or SB431542 (*SI Appendix*, Fig.S2E). Indeed, inhibition of γ-secretase activity by DAPT abolished HES1 promoter activity in podocytes exposed to GH or TGF-β1 (Fig.2F). Notch activation in podocytes isolated from GH administered mice was also ameliorated in the presence of SB431542 or AG490 (Fig.2G&H). Together, these data suggest that GH activates Notch signaling in podocytes and it is mediated via TGF-β1.

**Figure 2.**
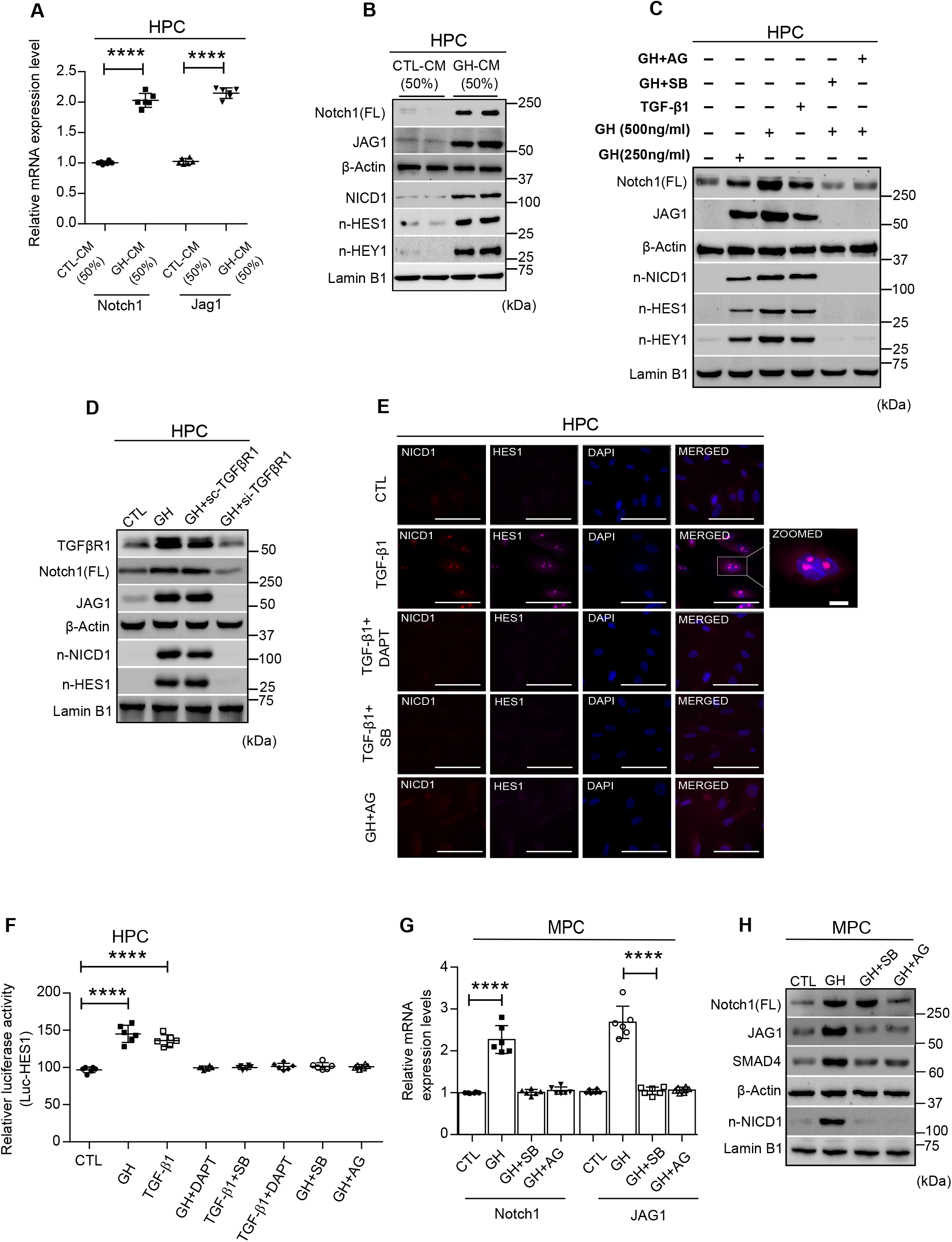
GH induces TGF-β/SMAD pathway mediated Notch signaling in podocytes. **(A)** qRT-PCR analysis showing the expression of Notch1 and Jag1 in HPC treated with or without conditioned medium (CM) of 50% from CTL and 50% GH treated podocytes for 48h. Mean±SD. (n=6). ****p<0.0001 by student t-test. mRNA levels were normalized to β-Actin and presented as fold-change on the y-axis. **(B)** Immunoblotting analysis from HPC treated with or without CM of 50% from CTL and 50% GH treated podocytes for 48hr. (n=3). **(C)** Immunoblotting analysis from HPC treated with or without GH (250 and 500ng/ml), TGFβ-1 (5ng/ml), GH (500ng/ml) + SB431542 (100nM/ml) and GH (500ng/ml) + AG490 (10μM/ml) for 48hr. (n=3). SB431542 is an inhibitor for TGFBR1 and AG490 is an inhibitor for GHR. n-NICD1 (nuclear-NICD1), n-HES1 (nuclear HES1) and n-HEY1(nuclear HEY1). **(D)** HPC cells transfected with specific siRNA targeting TGFBR1 or scramble RNA (Scr-RNA) were subjected to immunoblotting. (n=3). **(E)** Immunofluorescence for nuclear co-localization of NICD1 (red color), HES1 (purple color) and counterstained with DAPI (blue color) in HPC from CTL vs treatments for 48hr. Magnification x630. Scale bar=2Oμm. (n=3). **(F)** HES1 reporter activity was measured in HPC from CTL vs treatment for 48h. Mean±SD. (n=6). ****p<0.0001 by Student’s t-test. DAPT (5μg/ml). **(G)** qRT-PCR analysis of Notch1 and JAG1 expression in MPC from CTL or GH (1.5mg/kg bw), GH+ SB431542 (1mg/kg bw) and GH+ AG490 (1mg/kg bw) administered mice (each group, n=6). Mean±SD. (n=6). ****p<0.0001 by Student’s t-test. Each dot represents the average value of three independent experiments from a single mouse. **(H)** Immunoblotting analysis for MPC from CTL vs treatment group of mice. (n=3). β-Actin and Lamin-B1 served as an internal control.

### 3. Both GH and TGF-β1 induce cell cycle re-entry of quiescent podocytes in a Notch1 dependent manner

In healthy mice and human; mature podocytes are in quiescent stage (G0 phase), a prerequisite for their highly specialized functions [17]. Mature podocytes are terminally differentiated and express high levels of cyclin-dependent kinase (CDK) inhibitors suggesting that these cells lack the ability to renew during adult life [18]. Since Notch signaling was shown to induce proliferation of embryonic stem cells and cell cycle re-entry of terminally differentiated cells [19, 20], we assessed whether activated Notch signaling induces cell cycle re-entry of podocytes in our experimental setting. When podocytes were stained for α-tubulin, we found that 27±10% of GH or TGF-β1 treated cells were in anaphase as suggested by microtubule formation (Fig.3A&C), whereas DAPT prevented GH-induced cell cycle progression (Fig.3A&C). Live-cell imaging confirmed that GH-treated podocytes were accumulated in anaphase (Suppl. Movie1). The morphological screening of podocytes by phalloidin staining revealed 24±7% of GH or TGF-β1 treated cells were bi-nucleated and hypertrophic (Fig.3A&C). This aberrant phenotype was not observed in podocytes treated with AG490 and SB431542 (Fig.3A&C). Interestingly, inhibition of Notch by DAPT mitigated GH induced podocyte binucleation vis-a-vis hypertrophy (Fig.3A&C).

**Figure 3.**
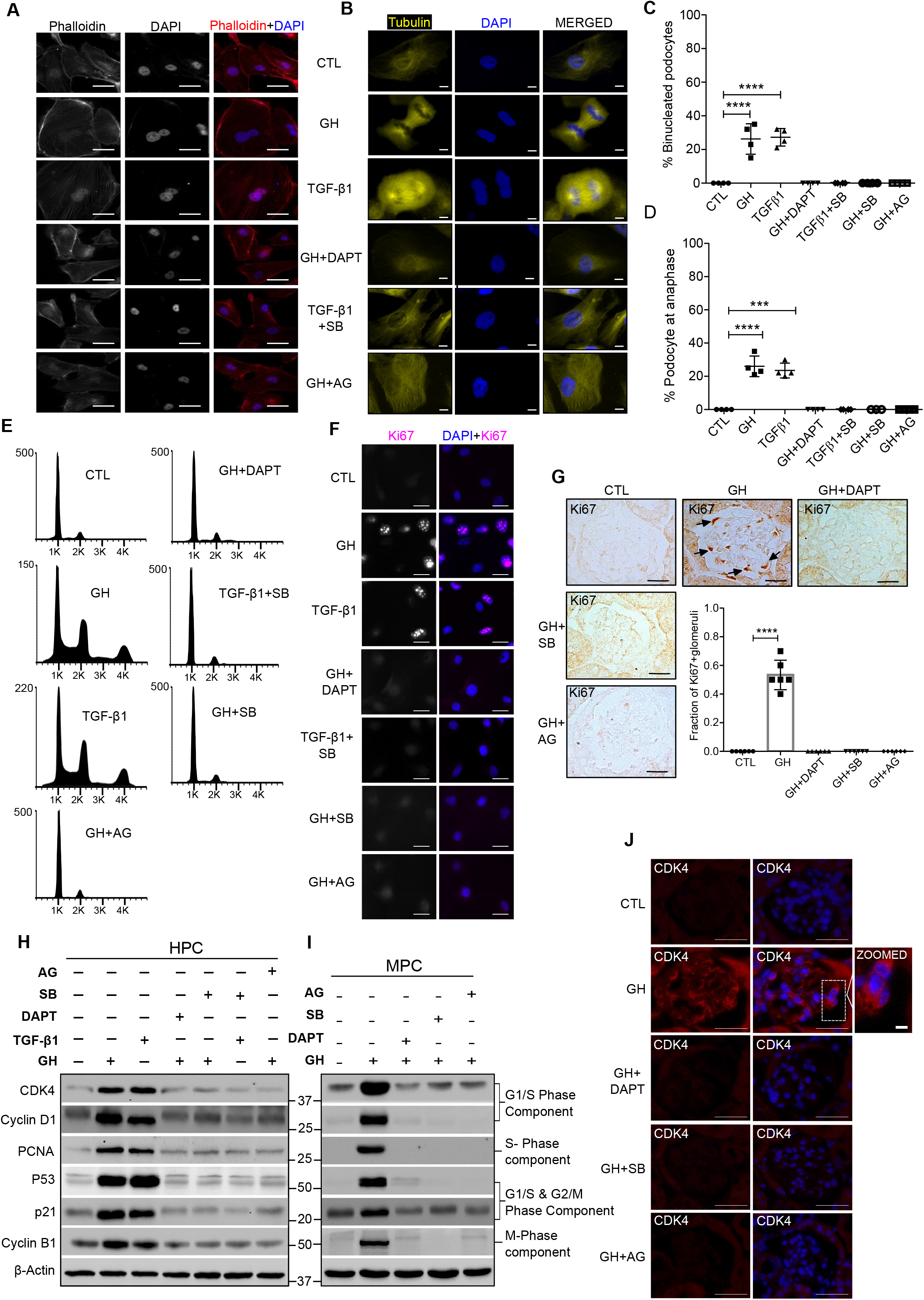
GH stimulates cell cycle re-entry and binucleation in differentiated podocytes. **(A&B)** Representative images of F-actin (red color), α-Tubulin (Yellow color), and counter-stained with DAPI (blue color) staining in HPC from CTL vs treatment for 48hr. Magnification x630. Scale bar=20 μm. (n=3). **(C)** The representative graph showed the percentage of binucleated HPC from CTL vs treatment for 48hr. Mean±SD. (n=4). ****p<0.0001 by student t-test. Each dot representing the average value of fifty cells. **(D)** Representative graph showed the percentage of HPC at anaphase from indicated CTL vs treatment for 48hr. Mean±SD. (n=4). ****p<0.0001 by student t-test. Each dot representing the average value of fifty cells. **(E)** Cell cycle phases of HPC from CTL vs treatment for 48hr. (n=4). **(F)** Immunofluorescence for the Ki67 (red color) and counterstained with DAPI (blue color) in HPC from CTL vs treatment for 48hr. Magnification x630. Scale bar=20μm. (n=3). **(G)** Representative images for anti-Ki67 expression by DAB staining in mice glomerular sections from CTL vs treatment group and graph represent the quantification of Ki67^+^ glomeruli. Black arrow indicates specific expression of Ki67 in podocytes. Magnification x630. Scale bar:20μm. (n=3). **(H&I)** Immunoblotting analysis from HPC (48hr treatment) and MPC from CTL vs treatment group. (n=3). **(J)** Representative images for CDK4 (red color) and counterstained with DAPI (blue color) in glomeruli from CTL vs treatment group. Magnification x630. Scale bar:20 μm. (n=3). White arrowhead indicates specific expression of CDK4 in podocyte. β-Actin served as internal control.

To further ascertain the activation of cell cycle events with GH or TGF-β1 we performed flow cytometric analysis (Fig.3E). Flow cytometry data revealed that ~40% & 31% of GH treated podocytes were in S and G2/M phase, respectively. Similarly, ~35%&28% of TGF-β1 treated podocytes were also accumulated in the S and G2/M phase, respectively (Fig.3E). As anticipated, AG490 and SB431542 abrogated cell cycle progression (Fig.3E). We observed elevated Ki67 expression, which is strongly associated with cell proliferation in GH treated podocytes both *in vitro* and *in vivo* (Fig.3F&G). Despite podocytes displaying proliferative phenotype when exposed to GH and TGF-β1, they also showed bi-nucleation suggesting only successful karyokinesis, but not cytokinesis. Therefore, in addition to proliferating markers (PCNA), we analyzed the expression of cell cycle regulators and checkpoints. Interestingly, in addition to cell cycle activators (CDK4 & CyclinD1), we also observed elevated expression of both G1/S and G2/M checkpoints, implying a complex two-tier regulation of cell-cycle events in podocytes exposed to GH or TGF-β1 (Fig.3H-J). Inhibition of GHR or TGF-βR1 mitigated Ki67 expression and also attenuated cell-cycle regulators in both human and mouse podocytes (Fig.3H-J). Strikingly, inhibition of Notch activation by DAPT abrogated GH-induced proliferating markers and in turn activation of cell cycle events (Fig.3F-J). Together, these data reveal that podocytes overcome quiescent stage and re-enter the cell cycle during stimuli such as exposure to GH or TGF-β1 in a Notch1 dependent manner.

### 4. Cytokinesis failure induces apoptosis in GH or TGF-β1 treated podocytes

Incomplete cytokinesis could be one of the predominant possibilities for binucleation that might arise as a result of aberration in contractile ring assembly or ingression phase of cytokinesis. RhoA, a member of the Rho GTPase family, is essential for cytokinesis via acting at the mid-body during cleavage furrow ingression and successful generation of two daughter cells [21, 22]. Elevated expression of RhoA was observed in podocytes either treated with GH or TGF-β1 and also during ectopic expression of NICD1 (Fig.4A-C). On the other hand, inhibition of Notch reduced GH or TGF-β1 induced RhoA expression (Fig.4A-B). Interestingly, abnormal localization of RhoA expression (away from contractile ring) was observed in podocytes treated with GH (Fig.4D). Although GH elicited cell-cycle entry of quiescent podocytes, these cells fail to accomplish successful cell division.

**Figure 4.**
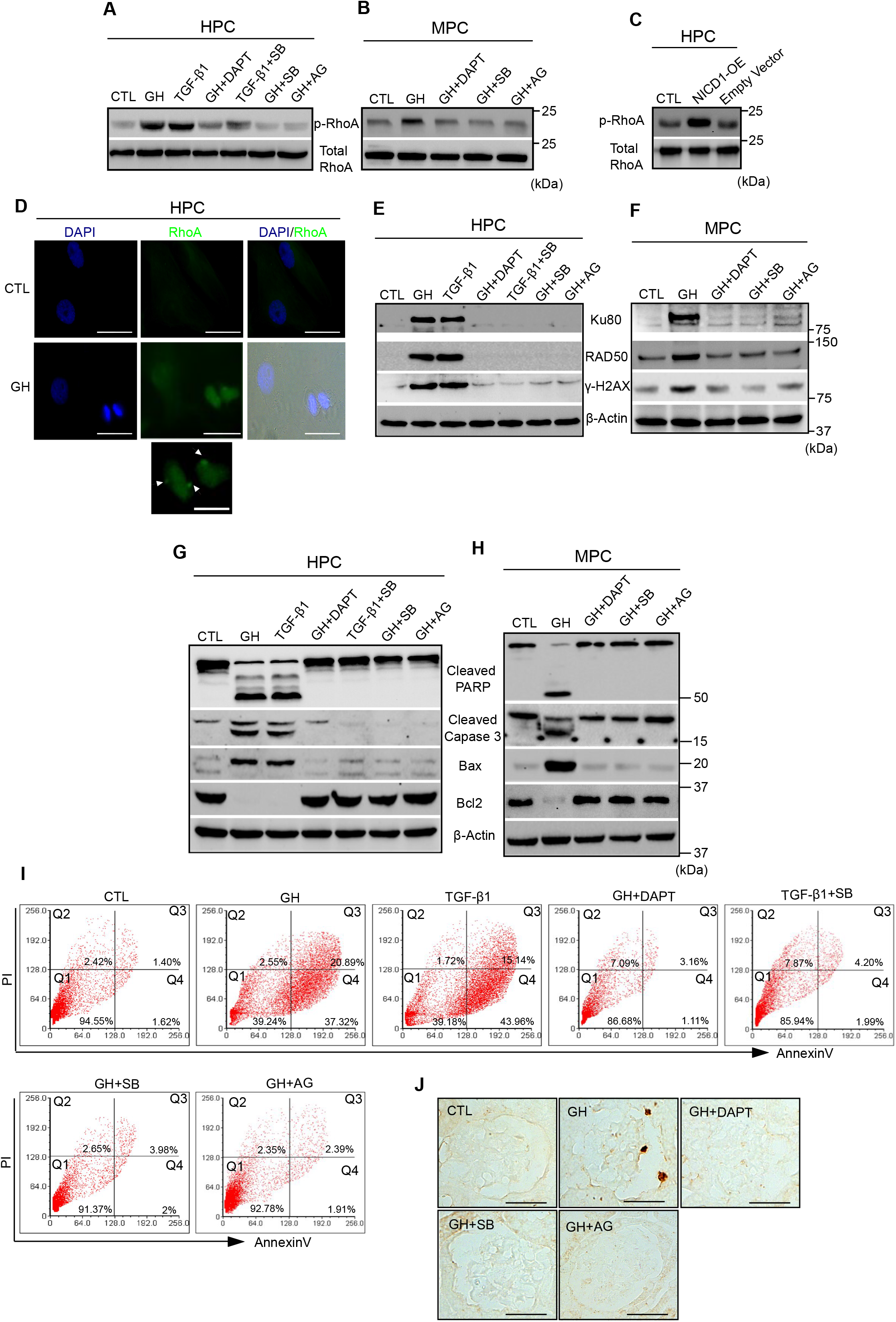
GH induced TGF-β leads to podocyte DNA damage and apoptosis. **(A&B)** Immunoblotting analysis for indicated genes expression in HPC (48hr treatment) and MPC from CTL vs treatment group. (n=3). **(C)** Immunoblotting analysis from HPC under ectopic expression of NICD1 (NICD1-OE). Empty vector denotes negative control. (n=3). **(D)** Immunofluo-rescence for the RhoA (green color) and counterstained with DAPI (blue color) in HPC from CTL vs treatment for 48hr. Magnification x630. Scale bar=20μm. (n=3). White arrowhead indicates the dislocalization of RhoA from midbody. **(E-H)** Immunoblotting analysis from HPC and MPC from CTL vs treatment group. (n=3). **(I)** HPC from CTL vs treatment for 48hr, stained with FITC AnnexinV and PI, and analyzed by flow cytometry. (n=4). The values of the representative histograms indicate the percentage of podocytes, in the lower left quadrant (Live cells), lower right quadrant (early apoptosis), upper right quadrant (late apoptosis) and upper left quadrant (necrotic cells). **(J)** Representative TUNEL (Terminal deoxynucleotidyl transferase dUTP nick end labeling) staining by DAB in glomerular sections from CTL vs treatment mice group. Magnification x630, Scale bar=20μm.(n=3). β-Actin served as internal control.

Often, failure of cell cycle progression is accompanied by induction of cell death. Therefore, we next investigated markers of DNA damage and apoptosis in podocytes exposed to GH or TGF-β1. Elevated expression of γ-H2X (a marker for DNA double-strand break), Ku80, and Rad50 (markers for double-strand repair) was observed in GH and TGF-β1 treated human podocytes and also in podocytes isolated from mice administered with GH (Fig.4E&F). Activated PARP & Caspase-3, induction of Bax (pro-apoptotic markers), and suppression of Bcl2 was observed in podocytes exposed to GH or TGF-β1 (Fig.4G&H, *SI Appendix*, Fig.S3A&B). Interestingly, the expression of GH or TGF-β1 induced DNA damage (Fig. 4E&F) and apoptotic (Fig.4G&H) markers were ameliorated by DAPT and AG490 or SB431542.

The majority of GH or TGF-β1 treated podocytes are early apoptotic (40±5% vs late 15±7%) (Fig.4I). Caspase 3 staining also revealed podocyte apoptosis with characteristic blebbing of the cell body (*SI Appendix*, Fig.S3C). However, DAPT, SB431542, and AG490 prevented podocyte apoptosis (Fig.4I & *SI Appendix*, S3C). Similarly, TUNEL staining also showed increased podocyte apoptosis in GH treated mice whereas DAPT, SB431542, and AG490 treatment ameliorated GH-induced apoptosis (Fig.4J). Our data suggest that GH or TGF-β1 treatment despite inducing the activation of mitosis, evoked cell death, a phenomenon is known as mitotic catastrophe [23].

### 5. Blocking of TGFB R1 or Notch1 signaling abrogates GH-induced podocytopathy and proteinuria

GH administered mice showed increased urinary albumin creatinine ratio (UACR) and proteinuria, and a decline in glomerular filtration rate (GFR) (Fig.5A-D). Elevated TGBR1 and CTGF and decreased BMP-7 expression was noticed in podocytes isolated from GH-treated mice (Fig.5E&F). CTGF is a TGF-β1 target whereas BMP-7 is an antagonist of TGF-β1. Furthermore, we also noticed the activation of canonical SMAD signaling in podocytes isolated from GH-treated mice (Fig.5F). As expected, blocking GHR (by AG490) or TGBR1 (by SB431542) prevented activation of SMAD signaling in GH administered mice (Fig.5E&F). Since GH treated podocytes showed bi-nucleation and consequent apoptosis *in vitro*, to ascertain *in vivo* confirmation, we counted the average number of podocytes per glomerulus in mice administered with GH. The number of podocytes (WT1 positive) in GH treated mice decreased significantly (P<0.005) compared with mice naive to GH treatment (Fig.5G, left & right panel). Blunting the Notch activation or inhibition of GHR/TGFBR1 mitigated GH-induced podocyte loss. Expression of slit-diaphragm proteins (podocin and ZO-1) was decreased in podocytes from GH treated mice (*SI Appendix*, Fig.S4A). Since podocyte loss and damage to the slit-diaphragm eventually manifest in glomerulosclerosis in addition to proteinuria [24, 25], we measured the histopathological changes by PAS, MT, H&E staining, and TEM imaging. Severe glomerulosclerosis (PAS and MT staining), and altered morphology (H&E staining) of the nephron was observed in GH treated mice, whereas TEM images revealed podocyte foot process effacement and thickening of the glomerular basement membrane (Fig.5H). Blunting of GH or TGF-β1 action or blocking Notch activation prevented GH-induced renal manifestations (Fig.5H). Suppression of GH or TGF-β1 action or preventing Notch signaling, preserved the expression of slit-diaphragm proteins (podocin & Z01) and prevented proteinuria. Blunting GH or TGF-β1 action also ameliorated TGF-β1 loss into the urine (*SI Appendix*, Fig.S4B). Together, our data demonstrate that GH’s role in the pathogenesis of nephropathy is mediated by TGF-β1.

**Figure 5.**
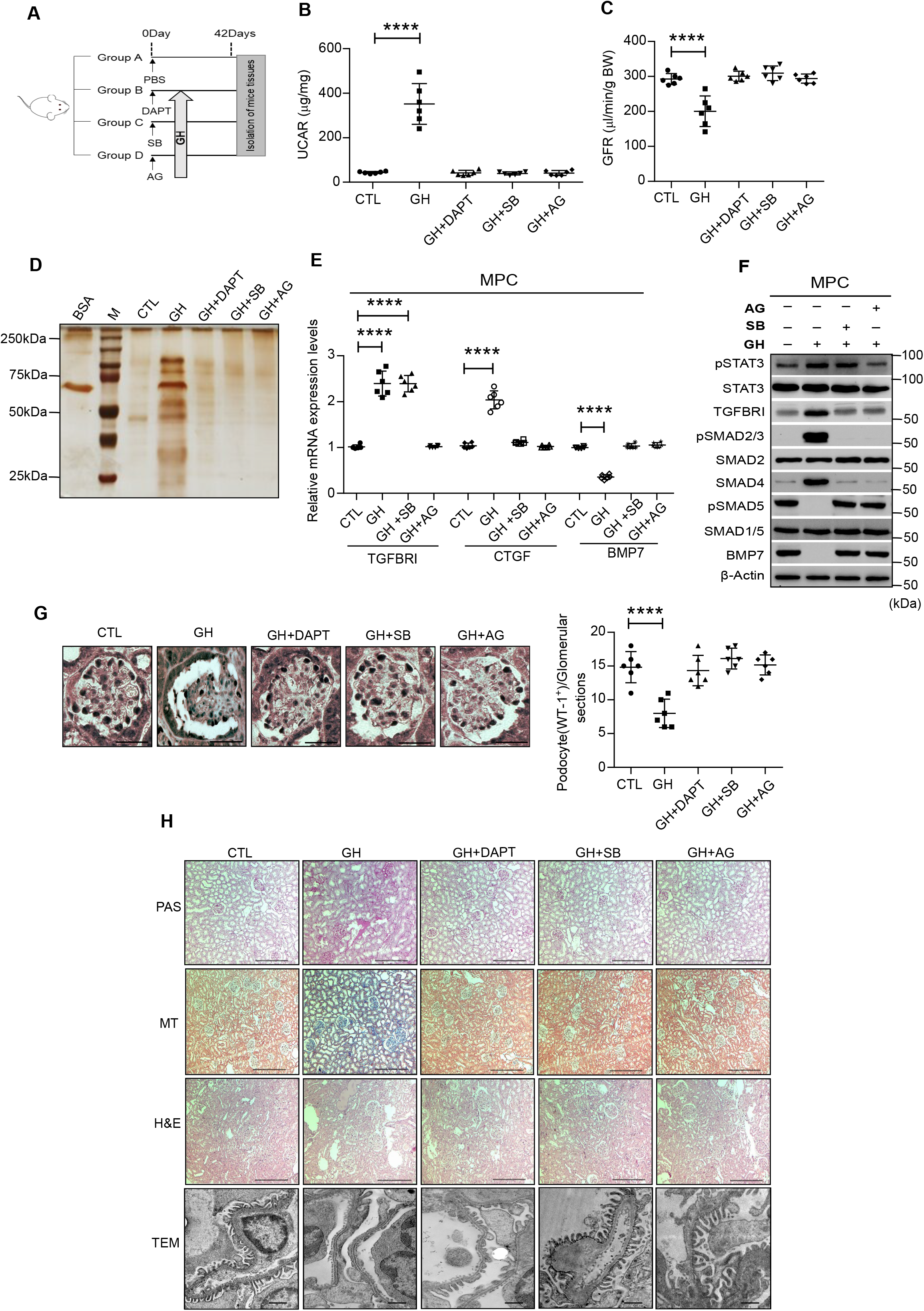
Blockade of GHR and TGFβRI protects mice from GH-induced proteinuria. **(A)** Schematic presentation of mouse experimentation. **(B)** Urinary albumin creatinine ratio (UACR) and **(C)** Glomerular Filtration Rate (GFR) were estimated in CTL vs treatment group of mice. Mean±SD. (n=6). ****p<0.0001 by Student’s t test. Each dot represents the average value of three independent experiment from a single mouse. **(D)** Silver staining was performed to the urine samples from CTL vs treatment group of mice. n=3. BSA; Bovine Serum Albumin, M; protein standard marker. **(E)** qRT-PCR analysis showing the expression of indicated genes in MPC from CTL vs treatment group of mice. Mean±SD. (n=6). ****p<0.0001 by Student t-test. Each dot represents the average value of three independent experiment from a single mouse. **(F)** Immunoblotting analysis for MPC from CTL vs treatment group of mice. (n=3). **(G)** *Left panel:* Representative images of immunohistochemical staining for anti-WT1 (podocytes) by DAB in the glomerulus sections from CTL vs treatment group of mice. Magnification x630, Scale bar=20μm.(n=3). *Right panel:* Average number of WT1+ cells in the glomerulus was quantified in mice from CTL vs treatment group with the help of ImageJ (NIH). Mean±SD. (n=6). ****p<0.0001 by student’s t-test. Each dot represents the average value of three independent experiments from a single mouse. **(H)** Representative image of PAS (Periodic acid-Schiff), MT (Masson’s trichrome), H&E (Hematoxylin and Eosin) staining in kidney tissue, and TEM (Transmission Electron Microscopy) analysis in podocytes from CTL vs treatment group of mice. Magnification x100. Scale bar= 100μm, TEM scale bar 1μm. β-Actin served as internal control.

### 6. Hyperactivated Notch signaling and binucleated podocytes in patients with diabetic nephropathy

We evaluated the extent of NICD1 expression and binucleation of podocytes in subjects with diabetic nephropathy (DN). Kidney biopsy sections from diabetics showed increased TGF-β1 and active Notch (NICD1) expression (Fig.6A). Furthermore, we observed both binucleated podocytes and also detached podocytes localized to urinary space in renal sections from people with DN (Fig.6B&C). As anticipated, glomerulosclerosis was observed in kidney sections from people with DN (Fig.6D). We have also observed elevated urinary TGF-β1 levels from these subjects with DN (Fig.6E&F). As expected, these diabetics showed severe proteinuria (Fig.6G). Interestingly, Nephroseq (https://www.nephroseq.org) analysis revealed co-expression of TGFBR1, Notch signaling components (HES1), cell proliferating markers (Ki67 & PCNA), cell cycle regulator (TP53), and regulator of cytokinesis (RhoA) in human diabetes glomerulus dataset (Woroniecka) (Fig.6H). Schematic illustration of GH action on podocyte cell cycle re-entry and death via TGF-β1 mediated Notch1 activation (Fig.6I). All these data confirm that people with DN have elevated functional Notch signaling in their glomeruli, podocytes with aberrant cell-cycle entry, and enhanced podocyte injury markers.

**Figure 6.**
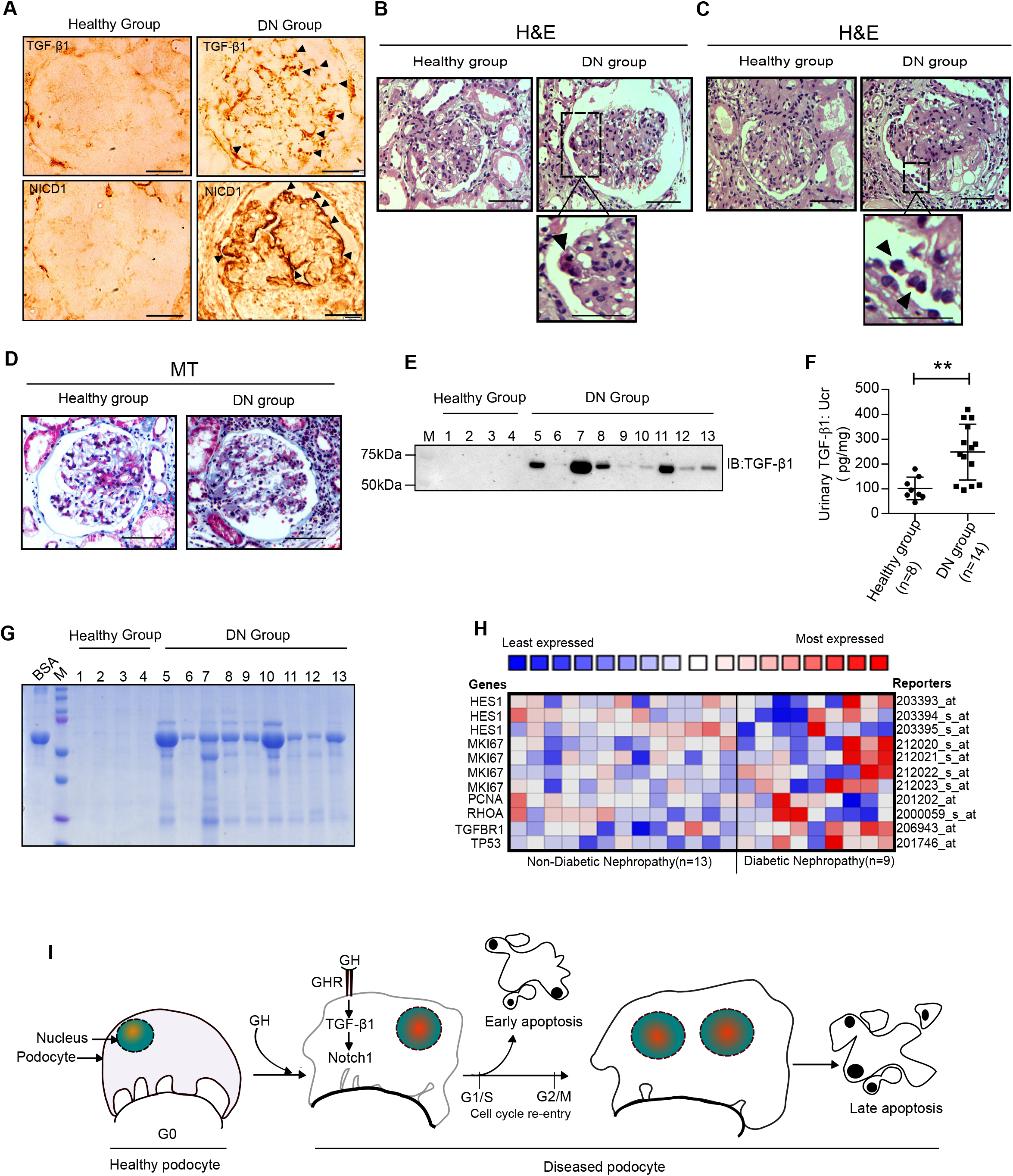
Elevated TGF-β1 signaling and proteinuria correlated in people with DN. **(A)** Representative images of immunohistochemical staining for TGFβ-1 and NICD1 by DAB in the glomerulus sections from healthy (n=8) and DN group (n=14). Magnification x630. Scale bar=20μm. (n=3). **(B&C)** Representative image of H&E stain in glomerular sections from healthy and DN group. Black arrowhead indicates bi-nucleated and detached podocyte. Magnification x630. Scale bar=20μm. Zoomed picture emphasizes a bi-nucleated and detached podocyte. **(D)** Representative image of MT stain in glomerular sections from healthy and DN group. Magnification x630. Scale bar=20μm. **(E)** Immunoblotting analysis for TGF-β1 in the urine samples from healthy (n=4) and DN group (n=9). **(F)** Quantification of TGF-β1 in the urine samples from healthy (n=8) and DN (n=14). Mean±SD. **p<0.001 by student’s t-test. **(G)** Urine samples from healthy (n=4) and DN (n=9) were resolved on SDS-PAGE and stained with Coomassie Blue. BSA= Bovine serum albumin. M=protein marker. **(H)** Nephroseq (www.nephroseq.org) analysis comparing HES1, MIKI67, PCNA, RHOA, TGFBR1, and TP53 expression levels in non-diabetic (n=13) versus diabetic nephropathy (n=9) (Woroniecka Diabetes glom, nephroseq.org). Data indicate that the expression of these genes increased >1.5-fold in the DN. **(I)** Schematic illustration of GH action on podocyte cell cycle entry and apoptosis via TGF-β1 mediated Notch1 activation.

## Discussion

The current investigation reveals a novel mechanism for GH action on glomerular podocytes and in the pathogenesis of DN. The major findings of this study are that GH induces TGF-β1 and the canonical TGF-β1/SMAD signaling in podocytes. GH and TGF-β1 activate Notch signaling, which is implicated in the cell cycle re-entry of podocytes. However, these activated podocytes fail to complete the mitotic cycle and as a consequence, binucleated podocytes are accumulated. Indeed, renal biopsies from patients with DN also revealed binucleated podocytes. Podocytes exposed to GH or TGF-β1 fail to accomplish mitosis due to cytokinesis failure and susceptible to cell death. Importantly, inhibition of GHR and TGFBR1 successfully protected mouse podocytes from GH and TGF-β1 induced Notch activation, cell cycle re-entry, and apoptosis. Furthermore, we observed podocyte loss, glomerulosclerosis, and proteinuria in GH treated mice, a common pathological feature associated with DN. The current study demonstrates that GH’s role in the pathogenesis of nephropathy is at least partly mediated by the TGF-β1/SMAD signaling. Interestingly, inhibition of Notch with DAPT significantly ameliorated both GH and TGF-β1 induced podocyte injury and apoptosis.

The first important observation from our study was that GH induces TGF-β1 expression in podocytes. Elevated GH levels are implicated in the early renal hypertrophy, depletion of podocytes, and proteinuria [26, 27]. However, it was not clear whether this causal role of the GH in the pathogenesis of nephropathy is due to direct actions of GH on the podocytes or via GH’s effector molecules. Among several hosts of mediators in the diabetic milieu, TGF-β1 has emerged to have a key role in the development of morphologic manifestations and clinical characteristics of diabetic nephropathy [22, 28-30]. Inhibition of TGF-β1 or ablation of SMAD3 (SMAD^-/-^) showed promising protection from glomerulosclerosis and renal dysfunction [13, 31]. Despite knowing the fact that activation of TGF-β/SMAD signaling is crucial in most kidney diseases, the stimuli that activate this pathway in the podocyte remain unclear. Previous studies proposed that high glucose and angiotensin II induces TGF-β1 expression in glomerular cells [30, 32]. Although, elevated GH levels and overactivity of the GH/GHR axis are implicated in renal manifestations and CKD [33], the temporal association between GH and TGF-β was unclear. It is noteworthy that GH-induced mild glomerulosclerosis and interstitial fibrosis in diabetic Sprague-Dawley rats is associated with an elevation in TGF-β1 levels [12] and suppressing JAK2, an immediate downstream target of GH reduced TGF-β mRNA expression [13]. In the present study, we establish that GH stimulates TGF-β1 in podocytes and to the best of our knowledge this is the first study to demonstrate that GH induces TGF-β1 expression.

Another key finding that emerged from our study is that both GH and TGF-β1 trigger cell cycle re-entry of podocytes by Notch activation. Notch is a highly conserved juxtracrine signaling cascade, which transduces short-range signals between neighboring cells. This pathway comprises 4 transmembrane receptors (Notch 1-4) and 5 ligands (Jagged 1&2; Deltalike 1, 3, & 4). Binding of ligand to Notch receptor results in shedding of both the Notch extracellular domain by ADAM protease and cleavage of the Notch intracellular domain (NICD) by the γ-secretase complex. Subsequent nuclear translocation of NICD activates expression of target genes such as Hes1 and Hey1. We had previously demonstrated that GH induces Notch activation [9]. Since TGF-β1 is a powerful Notch activator, we investigated whether the observed Notch activation in GH treated conditions could be due to TGF-β1 secreted under GH stimuli. Although active Notch signaling is required till the stage of S-shaped body formation during glomerulogenesis, it is almost undetectable in healthy adult glomeruli. Indeed, the down-regulation of the Notch pathway is required for renal progenitors to differentiate towards podocyte lineage [15]. Mature podocytes exit from the cell cycle which is evidenced by reduced expression of proliferating markers (eg: Ki67, PCNA, and CyclinB1) and increased expression of cell-cycle inhibitors p57and p27 [34]. The persistent expression of p57 and p27 enable the mature podocytes remain quiescent [35, 36]. Our data reveal that GH or TGF-β1 stimulated quiescent podocytes re-enter the cell-cycle and progressed to the S-and G2/M phase, and the progression of cell cycle events is concomitant with activation of Notch signaling. It was reported that Notch activation stimulates cell-division in renal progenitors whereas in differentiated podocytes it helps cells overcome the G2/M checkpoint [15]. Increased Notch activity was observed in podocytes of patients with glomerular disorders, and blunting Notch activity ameliorated glomerulosclerosis, prevented podocyte death during the initial phases of glomerular injury, and proteinuria [16].

A significant observation from this study is that podocyte exposed to GH become binucleated and hypertrophic. Normally, post-mitotic cells do not re-enter the cell cycle when exposed to growth signals. In the case of podocytes, we observed an increase in cell size with GH treatment and it caused increased kidney mass in these animals. Stressed podocytes were considered to re-enter the cell cycle and are arrested at G2/M restriction point by CDK inhibitors, and become hypertrophic [37]. Multi-nucleation of podocytes contributes to the increased cell size. The hypertrophic phenotype of podocytes appears to be transient as podocytes with cytokinesis failure and aneuploidy are susceptible to cell death [23]. Podocyte depletion has been considered as a hallmark of both primary and secondary forms of glomerulosclerosis [38-40]. Decreased podocyte density strongly correlates with the severity of proteinuria and DN progression [41-43]. A large body of evidence suggests that podocytes undergo apoptosis, which is considered as a major form of podocyte loss that culminates in glomerular injury [44, 45]. EMT of podocytes is considered an alternative cause for podocyte loss [46]. We have noticed both early and late phase apoptosis in podocytes exposed to GH or TGF-β1. Cell cycle transition from G1 to S phase leads to extensive DNA damage that culminates in the early apoptosis of podocytes. Whereas late apoptosis suggests that the podocytes with DNA damage that overcome the G2/M checkpoint, eventually failed to complete cytokinesis. These cells accumulated at the G2/M phase of the cell cycle with increased DNA content per cell, and eventually undergo late apoptosis. Most of the podocytes in our experiments underwent early apoptosis in response to GH treatment, suggesting that these terminally differentiated podocytes are not sufficiently competitive to carry cell cycle events successfully despite mitogenic stimuli by GH or TGF-β1. Wu et al. reported that TGF-β at a lower concentration promotes podocyte differentiation whereas TGF-β levels beyond a critical threshold induce G2/M block and apoptosis [47]. Whereas GH plays a significant role in normal renal function and overactive GH signaling has been implicated in proteinuria in diabetes. Together these studies suggest in a dose-dependent manner GH and TGF-β specify podocyte fate.

Mature podocytes possess high cytoplasm to nucleus ratio and express highly organized myofibrils that normally prevent cell division. Disassembly of cytoskeletal filaments during mitogenic stimuli would adversely affect their function. Another interesting feature of differentiated podocytes is that they express a wide range of cell cycle proteins, which could be a prerequisite for executing mitotic catastrophe in response to stress signals such as GH and TGF. The mechanisms by which cell cycle re-entry causes cell death are not completely explained, but cytokinesis failure and abnormal Rho distribution as observed in our study could be one of the major reasons. Further studies are required to delineate the contribution of specific STATs in GH mediated TGF-β1 production, and whether the source of GH *in vivo* is more local than endocrine or opposite. Based on our observations, we propose that GH induces TGF-β1 expression and is a causative factor in the development of podocyte hypertrophy, podocyte injury, and consequent proteinuria during GH-induced kidney diseases. In summary, the present report establishes that GH induces TGF-β1 expression in podocytes and that some of the actions of GH on the podocytes are mediated through TGF-β1 in a both autocrine and paracrine manner. Our data provide a mechanistic link between GH and podocyte dysfunction in diseases like type I diabetes mellitus and acromegaly.

## Materials and Methods

### Antibodies and Reagents

The primary antibodies are as follows: anti-activated Notch1(ab8925), anti-pSTAT3 (ab76315), anti-STAT3(ab5073), TGF-βR1(ab31013), anti-HEY1(ab154077), p53 (ab26), RhoA(ab54835), Cyclin B1(ab72), p21(ab109520), α-Tubulin(ab7291), CDK4 (ab137675), Ki67(ab16667), CyclinD1(ab16663) and Laminin-B1(ab16048) were purchased from Abcam (Cambridge, MA). Anti-Notch1-Full-length(#3608), anti-cleaved-Notch1(#4147S), pSMAD2/3(#8828), anti-SMAD2(#5339), anti-pSMAD1/5(#9516), anti-SMAD5(#12534), anti-Cleaved-PARP(#5625), anti-Cleaved Caspase3(#9664), anti-BMP7(#4693), anti-BAX(#89477), anti-Bcl2(#15071) and anti-β-Actin(#4970) were purchased from Cell Signaling Technology (Danvers, MA). Anti-Ku80(NBP156408), anti-RAD50(NB100-147), anti-γ-H2AX(NB100-384), anti-Nephrin (NBP1-77303) and anti-PCNA(NBP500-106). The anti-TGF-β1(MAB240) from R&D Systems (Minneapolis, Minnesota). Anti-HES1(sc-166410) anti-WT-1(sc-393498) and anti-ZO-1(sc-33725) were obtained from Santa Cruz Biotechnology (Dallas, TX). Anti-JAG1(PAB807Hu01) was purchased from Cloud-clone (Houston, TX). Anti-Caspase3(9H19L2) and TGF-β1(BMS249-4) were purchased from Thermo Fischer Scientific (Waltham, Massachusetts). Mouse/Rabbit PolyDetector DAB HRP Brown-Bio SB (BSB020, Santa Barbara, CA). Phalloidin fluorescein isothiocyanate labelled (P5282) and glutaraldehyde solution (G5882) were obtained from Sigma chemicals (St. Louis, MO, USA). Precision Plus Protein Dual Color Standards (Bio-Rad, Hercules, CA), and ProLongTM Diamond Antifade Mountant (P36961) were purchased from Molecular Probes Life Technologies. DyLight 488 and DyLight 564, and Cy5-conjugated secondary antibody were purchased from Vector Laboratories (Burlingame, CA). Primers used in this study procured from Integrated DNA Technologies (Coralville, IA). All other reagents used were of analytical grade and obtained from Sigma chemicals (St. Louis, MO, USA).

### Experimental drugs

DAPT(D5942) was purchased from Sigma chemicals (St. Louis, MO, USA), TGF-βR1 inhibitor (SB431542) and JAK2 inhibitor (AG490) were purchased from Tocris Bioscience (1614-10MG). Recombinant TGF-β1 (#240-B-002) and human growth hormone Genotropin (Pfizer) procured from R&D Systems.

### Podocyte culture and experimentation

In this study, conditionally immortalized human podocytes (A gift from Prof. Moin Saleem, University of Bristol) cells were cultured essentially as described earlier [9]. Briefly, after 14 days of differentiation at 37°C podocytes were treated with or without GH, GH+DAPT, TGF-β1, TGF-β1+SB431542, GH+SB431542 and GH+AG490. Unless otherwise mentioned all the experimental conditions for podocyte cells were given for 48 hr. The cell lysate was prepared for RNA isolation, immunoblotting and Enzyme Immunosorbent Assay (ELISA). For immunofluorescence, cells were cultured on coverslips, followed by treatment as mentioned above, subsequent fixation with paraformaldehyde (4%), and blocking with PBS containing normal BSA (5%) before incubation with primary antibodies. The next day, the samples were incubated with Alexa Fluor-conjugated secondary antibodies, and DAPI for nuclear stain, for 1hr at room temperature. Images were acquired using a laser scanning microscope (ZESSI, Germany). For F-actin staining in podocytes cells essentially as described earlier [9]. Briefly, HPC cells were incubated with Fluorescent phalloidin-TRITC conjugate (P1951) for 40 min at room temperature. Next counterstaining by DAPI (P36971), mounting and images were acquired using a Leica trinocular microscope or Apotome Axio Imager Z2 (Zeiss). Images were analyzed with LASX Industry Software and ImageJ. For Live-cell imaging, cells were grown on μ-Dish 35 mm (#81156) at 60% confluency treatment performed for 48hr and then imaged for 2hr on a Leica SP5 confocal laser scanning microscope with a HCX PL APO CS 63×, 1.40-NA oil-immersion lens.

### Animal and Tissues

All the experimental procedures for the animals were approved by the Institutional Animal Ethics Committee of the University of Hyderabad, India. 8-Week-old Swiss Webster male mice weighing nearly 30±5 g was used in this study. The mice were randomly assigned to five groups (6 mice per group): 1) control group (CTL), 2) GH-treated group, 3) GH+DAPT-treated group, 4) GH+SB431542 and 5) GH+AG490. Experimental mice received a single i.p. hGH (1.5 mg/kg/day), whereas control mice have received an equal volume of saline for 6 weeks. The inhibitor groups were received DAPT (10 mg/kg of body weight) per day prior to the GH treatment. After 6 weeks of the experimental period, the mice were placed in individual metabolic cages for collecting 24 hr urine to estimate albumin (#COD11573) and creatinine (#COD11502) levels as recommended by the manufacturers protocol (Biosystems, Barcelona, Spain). An aliquot of urine from mice was subjected to SDS-PAGE gel and silver staining was performed to compare the urinary protein profile for all the five groups. Further, we have also measured the GFR in these mice, as described previously [9]. Mouse podocytes were isolated from kidney of mice as described in earlier protocol [48]. Briefly, glomeruli were prepared by filtration of the cortex of kidney with mesh sieves, whose holes were 100, 76, and 54 μm in diameter, then the tissues left on the mesh sieve with 54 μm holes were collected and prepared for the RT-PCR and immunoblotting. For histological analysis, kidney cortex was fixed with 4% paraformaldehyde before embedding in paraffin. Paraffin-embedded tissues were sliced longitudinally into 3-4 μm thick sections, subjected to staining with Periodic-acid Schiff Base (PAS), Masson’s trichrome (MT) and Haematoxylin and Eosin (H&E) staining. Transmission electron microscopic (TEM) images were obtained for glomerular sections from all the experimental mice groups as described earlier [9].

### RNA extraction and Quantitative RT-PCR assay

The total RNA was prepared from HPC and MPC by using a TRIzol RNA isolation reagent (Thermo Scientific, Waltham, MA). Next, 1 μg of total RNA was reverse transcribed using the cDNA synthesis kit (PrimeScript 1st strand cDNA Synthesis). qRT-PCR analysis was performed by the QuantStudio 3 system (Applied Biosystem) with SYBR Green (Kappa Biosystem) Master Mix as mentioned in the following protocol: initial denaturation at 95°C for 3 min, followed by 35 cycles of three steps each at 95°C for 30s, 60°C for 30s, and 72°C for 30s. mRNA expression of each gene was normalized using the expression of β-actin.

### Western Blotting

Cytoplasmic extract for immunoblotting was prepared as described previously [9]. Briefly, Human podocytes and isolated mouse primary podocytes were lysed by RIPA buffer (Cell Signaling) containing protease inhibitor mixture (Sigma–Aldrich) and phosphatase inhibitor tablets (Roche), centrifuged and collection of supernatants. However, for nuclear extract protein sample preparation, pellet was resuspended with 20 mM HEPES (pH 7.9), 25 % Glycerol, 0.42 M NaCl, 0.2 mM EDTA, 1.5 mM MgCl_2_, 1 mM DTT, 0.2 mM PMSF) and vortex for 20 sec. Incubate the cell lysate for 25 min on ice and vortex every 10 min for 10 sec. Next, cell lysate was centrifuged for 12 min at 13,500 rpm at 4°C and collect the supernatant (Nuclear). The protein concentrations of cell and mouse podocyte lysates were determined using a bicinchoninic acid reagent (Thermo Scientific) using bovine serum albumin as a standard. Total 20 to 25 μg of protein samples were resolved by sodium dodecyl sulfate polyacrylamide electrophoresis (SDS-PAGE; Bio-Rad, Madrid, Spain) followed by Western blot. Immunoblot bands were visualized using a ChemiDoc™ XRS System (Bio-Rad).

### Enzyme-Linked Immunosorbent Assay

γ-Secretase activity was quantified as describe earlier [9]. Briefly, HPC cells were treated with or without GH, TGF-β1, GH+DAPT, TGF-β1+SB431542, GH+SB431542 and GH+AG490 for 48 hr and cleavage-dependent release of APH-1A measured at 450 nm by using a fluorescent microplate reader (Multiskan O Microplate Spectrophotometer, ThermoFisher Scientific). For TGF-β1 detection in condition media from HPC treated with different experimental conditions and in urine samples from mice and human were analyzed by TGF-β1 ELISA commercial kit (R&D Systems) according to the manufacturer’s protocol.

### Cellular DNA Flow Cytometric Analysis

The single-cell suspension of HPC (5×10^5^ to 1×10^6^ cells) from with or without GH, TGF-β1, GH+DAPT, TGF-β1+SB431542, GH+SB431542 and GH+AG490 for 48hrs were prepared in 300 μL PBS, fixed by cold 70% ethanol for 30 min at 4°C and then washed and resuspended in 300 μL PBS, followed by treatment with 3 μL RNase at 37°C for 30 min, chilled on ice, and 30 μL PI (propidium iodide; Roche) treatment in the dark at room temperature for 1hr. DNA contents were acquired by a S3e Cell Sorter flow cytometer (Bio-Rad) using FCS Express 7 program.

### Apoptosis analysis in podocytes

Apoptotic cell death was measured by an Alexa Fluor® 488 annexin V and propidium iodide (PI) apoptosis detection kit (ThermoFisher Scientific) according to the manufacturer’s protocol. Briefly, HPC cells were plated on 6 cm dishes at 1×10^5^ cells per dish with or without GH, TGF-β1, GH+DAPT, TGF-β1+ SB431542, GH+SB431542 and GH+AG490 for 48 hr. Next podocytes were harvested with the help of trypsin, washed with cold PBS twice, resuspended in binding buffer, and stained with FITC-Annexin V and PI in the dark at room temperature for 15 min. After incubation, binding buffer was added, and the podocytes were analyzed by a S3e Cell Sorter flow cytometer (Bio-Rad). Unstained cells and cells stained with FITC-Annexin V or PI alone were used as controls to set up compensation and quadrants in flow cytometry. The results were analyzed by the FCS Express 7 program.

### Reporter assay

HES1 promoter activity luciferase assay was performed as describe earlier [49]. Briefly, HPC cells were transfected with a pHES1(467)-Luc (procured from Addgene) and internal control expressing the *Renilla luciferase*, pRL-TK (Promega). HPC cells were transfected using Xfect polymer (DSS Takara Bio, New Delhi, India) as per manufacturer’s instructions. After 48 hr of transfection, cells were treated with or without GH, TGF-β1, GH+DAPT, TGF-β1+ SB431542, GH+SB431542 and GH+AG490 for 48 hr and washed twice with PBS and harvested in 100 μl of passive lysis buffer. After a brief freeze-thaw cycle, the insoluble debris were removed by centrifugation at 12,000g (4°C for 5 min), and 20 μl of supernatant was used for luciferase assay. The activity of the co-transfected luciferase reporter plasmid was used to normalize transfection efficiency.

SMAD4 cignal-GFP reporter assay was performed in podocyte cells according to the earlier published protocols with minor modifications [50]. Briefly, podocyte cells in a 96 well plate, 1000 cells/well were seeded prior to the day of transfection. Cells were transfected either with SMAD4-GFP, positive control and negative control vectors in triplicates. After transfection cells were left untreated in complete medium for 16hr. Then cells were treated with GH-CM, Anti-TGFβ1 antibody (#AB-100-NA), GH and TGFβ1. After 48hr of treatment, images were taken by using a Leica trinocular microscope. Fluorescence emission spectrum for these samples were acquired at 510-520nm by using a fluorescent microplate reader (Multiskan O Microplate Spectrophotometer, ThermoFisher Scientific).

For SMAD4 luciferase assay, Podocyte cells were seeded into a 6-well plate (2 × 10^5^ cells per well). The cells were then co-transfected with either 0.3 μg/well of Smad4 firefly luciferase reporter plasmid constructs (pLuc366 or pLuc207) or the control pGL3-Basic vector (Promega, San Luis Obispo, USA). The renilla luciferase plasmid was also co-transfected to correct for variations in transfection efficiency (45 ng/well). After incubating for 24hr, treatment was given for 48hr. Next cells were harvested from each experimental condition and the luciferase activity was measured using a fluorescent microplate reader (Multiskan O Microplate Spectrophotometer, ThermoFisher Scientific). Final activity was calculated as the ratio of firefly luciferase activity versus renilla luciferase activity units.

### Transfection of podocytes for knockdown and overexpression

HPC cells were trans-fected with siRNA as describe earlier [49]. Briefly, transfection was done using jetPEI reagent (Polyplus, Illkirch, France). HPC cells were seeded at 70-90% confluency in 6-well cell culture plates and transiently transfected with siRNA specific to TGFBR1 (#EHU051131) and its parental negative control siRNA by mixing with NaCl-jetPEI complexes. After 72 hr of transfection, cells were treated with or without GH, TGF-β1, GH+DAPT, TGF-β1+SB431542, GH+SB431542 and GH+AG490 for 48 hr. Next, cells were washed twice with PBS and lysed with RIPA buffer; the expression levels measured by western blotting as described above. The transient transfection of pT3-EF1aH NICD1; an overexpressing vector from Addgene (#86500) and its empty parental vector, pT3-EF1aH to the podocytes using Xfect polymer (DSS Takara Bio, New Delhi, India) as per manufacturer’s instructions. After 24 hr of transfection incubation, cells were harvested, and immunoblotting was performed.

### Statistical analysis

Statistical analysis methods are detailed in the figure legends.

### Ethics approval

The study was approved by the Institutional Review Board of Guntur Medical College and Government General Hospital, Guntur, India (#GMC/ IEC/120/2018) and adhered to the principles and the guidelines of the Helsinki Declaration. The animal experimental procedures were performed in adherence with the Institutional Animal Ethics Committee of the University of Hyderabad.

## Acknowledgments

The authors acknowledge Syed V Tahaseen for her help with human kidney specimens and urine samples. We thank Prof. Ram K. Menon (University of Michigan, Ann Arbor) for sharing microarray data. Authors thank Dr. Arun Kumar Kota for providing SMAD4-GFP signal reporter assay kit. The authors acknowledge Science and Engineering Research Board (SERB #EMR/2015/002076 and CRG/2019/005789) for providing funding to AKP.

## Authors contributions

RN, DM and AKP planned and designed the study. RN, DM and AKS performed the experiments. RN, AKP and PT evaluated the data. RN and KC performed the FACS experimentation and analysis. RN, DM and AKP wrote the manuscript. RN and AKP are the guarantor of this work and as such had full access to all the data in the study and takes responsibility for the integrity of the data and the accuracy of the data analysis.

