## Supplementary material for "Growth hormone induces mitotic catastrophe of podocytes and contributes to proteinuria": Fig.S

**Running title:** Growth hormone induces podocyte apoptosis

**Authors contributions:** RN, DM and AKP planned and designed the study. RN, DM and AKS performed the experiments. RN, AKP and PT evaluated the data. RN and KC performed the FACS experimentation and analysis. RN, DM and AKP wrote the manuscript. RN and AKP are the guarantor of this work and as such had full access to all the data in the study and takes responsibility for the integrity of the data and the accuracy of the data analysis.

[1]Authors contributed equally to the manuscript

**Figure S1. GH induces TGF-β1 in both HEPG2 cells and podocytes: (A)** qRT-PCR analysis showing the expression of TGF-β1 in HEPG2 cells treated with GH in concentration (100 to 500 ng/ml) and time (up to 48 h) dependent manner. mRNA levels were normalized to β-Actin and presented as fold-change. Mean±SD. (n=5). ****p<0.0001 by 1-way ANOVA post hoc Dunnett test. **(B)** Immunoblotting analysis for indicated genes from HEPG2 cells treated with GH in concentration (500 ng/ml) and time (up to 48h) dependent manner. (n=3). **(C)** Heatmap presenting differential expression of 14 genes in GH treated podocytes. Each vertical axis represents the gene rank with log2 fold change values. The color intensity of each row represents a mean centered log2 expression. Gene pattern (http://genepattern.broadinstitute.org/gp/) was used to generate this heat map. The original microarray data is available at GEO repository (GSE21327). **(D &E)** Immunoblotting analysis for indicated genes from HEPG2 and HPC cells treated with 10 to 50% conditioned media (GH-CM), GH-CM+SB (SB indicates SB431542, an inhibitor for TGFBR1) and GH-CM-Ab (Ab indicates ‘TGF-β1 neutralizing antibody’) for 48 h. (n=3).

**Figure S2. GH induces SMAD signaling and γ-secretase activity in podocytes both auto- and paracrine manner: (A)** Immunofluorescence for the nuclear co-localization of SMAD4 in GH treated human podocytes (HPC). Magnification x400. Scale bar=50 μm. (n=3). **(B)** SMAD4 luciferase activity in HPC treated with (48 h) or without GH. Mean±SD. (n=6). ****p<0.0001 by Student’s t-test. **(C)** SMAD-GFP signal reporter assay was performed in HEK293T cells treated with GH, CM-GH, and rTGF-β1. Un-transfected indicate there is no vector backbone. Negative CTL indicate empty vector backbone (Cignal SMAD4 without GFP). Positive CTL indicate Cignal SMAD4-GFP under treatment with 5 ng/ml TGF-β1. **(D)** Immunoblot for indicated genes in HPC exposed to conditioned media from GH treated podocytes (CM-GH) for 48 h. (n=3). **(E)** γ-secretase activity was measured in HPC from with or without GH(500ng/ml), TGF-β1 (5ng/ml), GH+DAPT(5μg/ml), TGF-β1 +SB431542 (100nM/ml), TGF-β1+DAPT, GH+SB431542 and GH+AG490 (10μM/ml). Mean±SD. (n=6). ****p<0.0001 by Student’s t-test.

**Figure S3. GH induced TGF-β leads to podocyte apoptosis: (A&B)** qRT-PCR analysis showing the expression of Bax and Bcl2 in HPC treated with GH or TGF-β for 48 h in the presence or absence CTL vs treatment for 48 h in the presence or absence of inhibitors for GHR and TGFBR1. mRNA levels were normalized to β-Actin and presented as fold-change on y-axis. Mean±SD. (n=5). ****p<0.0001 by Student’s t-test. **(C)** Immunofluorescence staining for Caspase 3 (red color) and counterstained with DAPI (blue color) in HPC from CTL vs treatment for 48 h. Magnification x630. Scale bar=20 μm. (n=6).

**Table1:** Table1. The list of qRT-PCR primers used in this study.

| | |
|---|---|
| TGFBR1(Human) | FP= TCAATTGTAAGCACATTGAAAGGG |
| | RP=TTCGCCCGGCAGATCTAAAC |
| TGFBR1(Mouse) | FP=AAGACAACTGCCAGCCCTTAG |
| | RP=TCATTTAGTGCCACACCCCA |
| TGF-β1(Human) | FP=GTTCAGGTACCGCTTCTCGG |
| | RP=CCTGATCGCCTCCCTTCATTT |
| TGF-β1(Mouse) | FP=AAATCAACGGGATCAGCCCC |
| | RP=CGCACACAGCAGTTCTTCTC |
| Notch1(Human) | FP=TGAATGGCGGGAAGTGTGAA |
| | RP=CACAGCTGCAGGCATAGTCT |
| Notch1(Mouse) | FP=AGACATGTAGGGCAGTCAGC |
| | RP=CCAGAGCTTACGTCATCCCA |
| HES1(Human) | FP=ATGACAGTGAAGCACCTCCG |
| | RP=GAGTGCGCACCTCGGTATTA |
| HES1(Mouse) | FP=TCCCACGGTCTGGGTCTTAT |
| | RP= GTGCTAAACCACTGACCCCT |
| JAG1(Human) | FP=CCTGTCCATGCAGAACGTGA |
| | RP=CGCGGGACTGATACTCCTT |
| JAG1(Mouse) | FP=GTTTCGCAGGAGGCCTGTTT |
| | RP=CTGGGTCAGCACCGAGAATG |
| BAX(Human) | FP=CTGACGGCAACTTCAACTGG |
| | RP=GCAGGGGGTTGATACCACG |
| Bcl2(Human) | FP=CGGGTTGTCGCCCTTTTCTA |
| | RP=TCACAGATCTGAGGGGGAGC |
| CTGF(Mouse) | FP=GCATCTCCACCCGAGTTACC |
| | RP=TAGGGGCAGAGGATGTACCTT |
| BMP7(Mouse) | FP=GTCTGCCAGGAAAGTGTCCA |
| | RP=CGAGGCTTGCGATTACTCCT |

**Movie legend:**

**Supply Movie1.** Live cell imaging of human podocytes treated with GH (500ng/ml for 48 h) and imaged for 2 h. We observed cells at anaphase stage and also cells under apoptotic phase.

**Supplymentary Figure1**

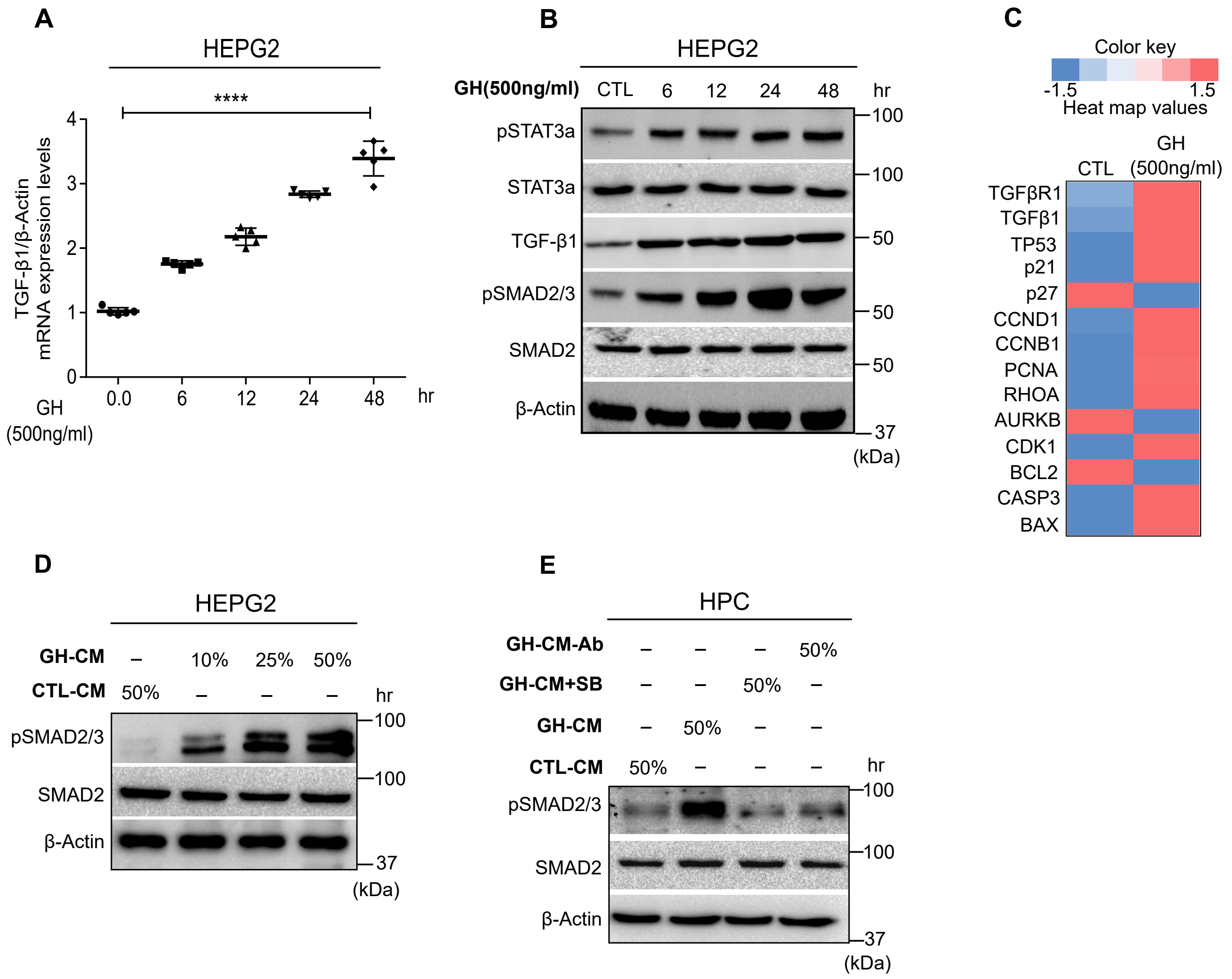

### Supplymentary Figure2

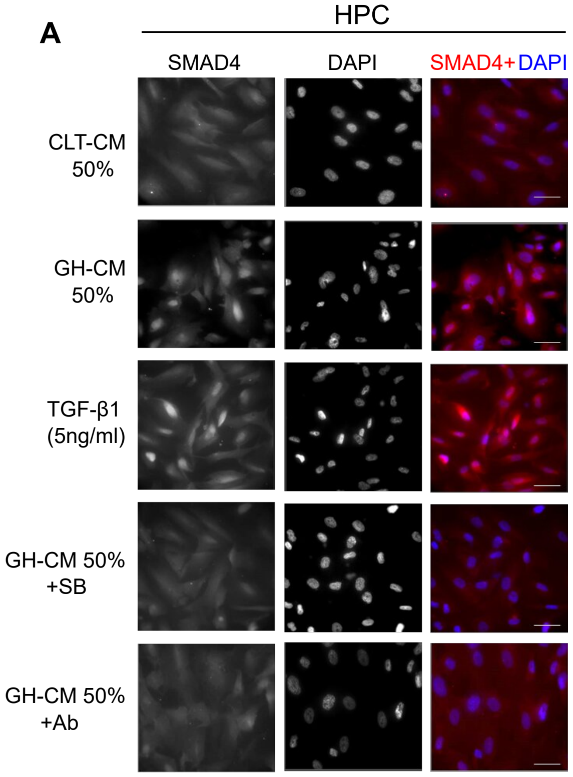


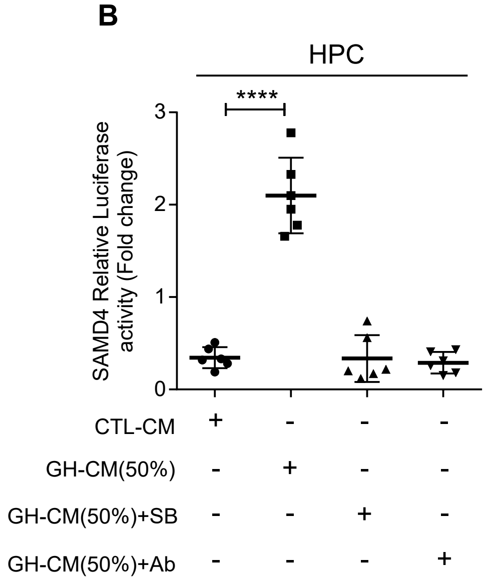


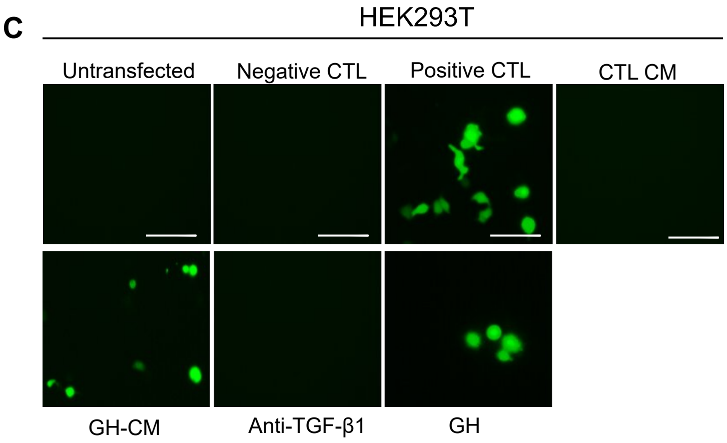


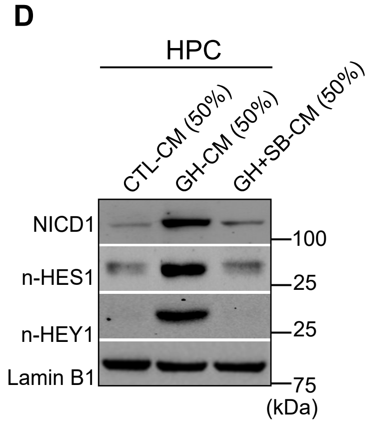


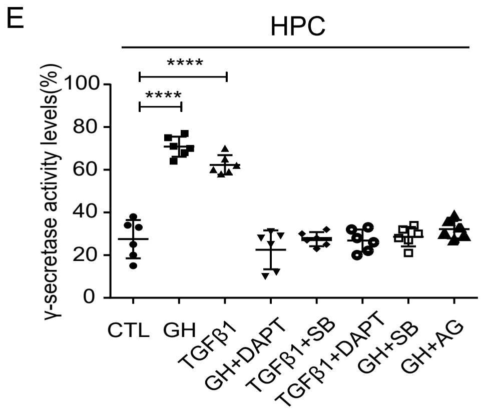

### Supplymentary Figure3

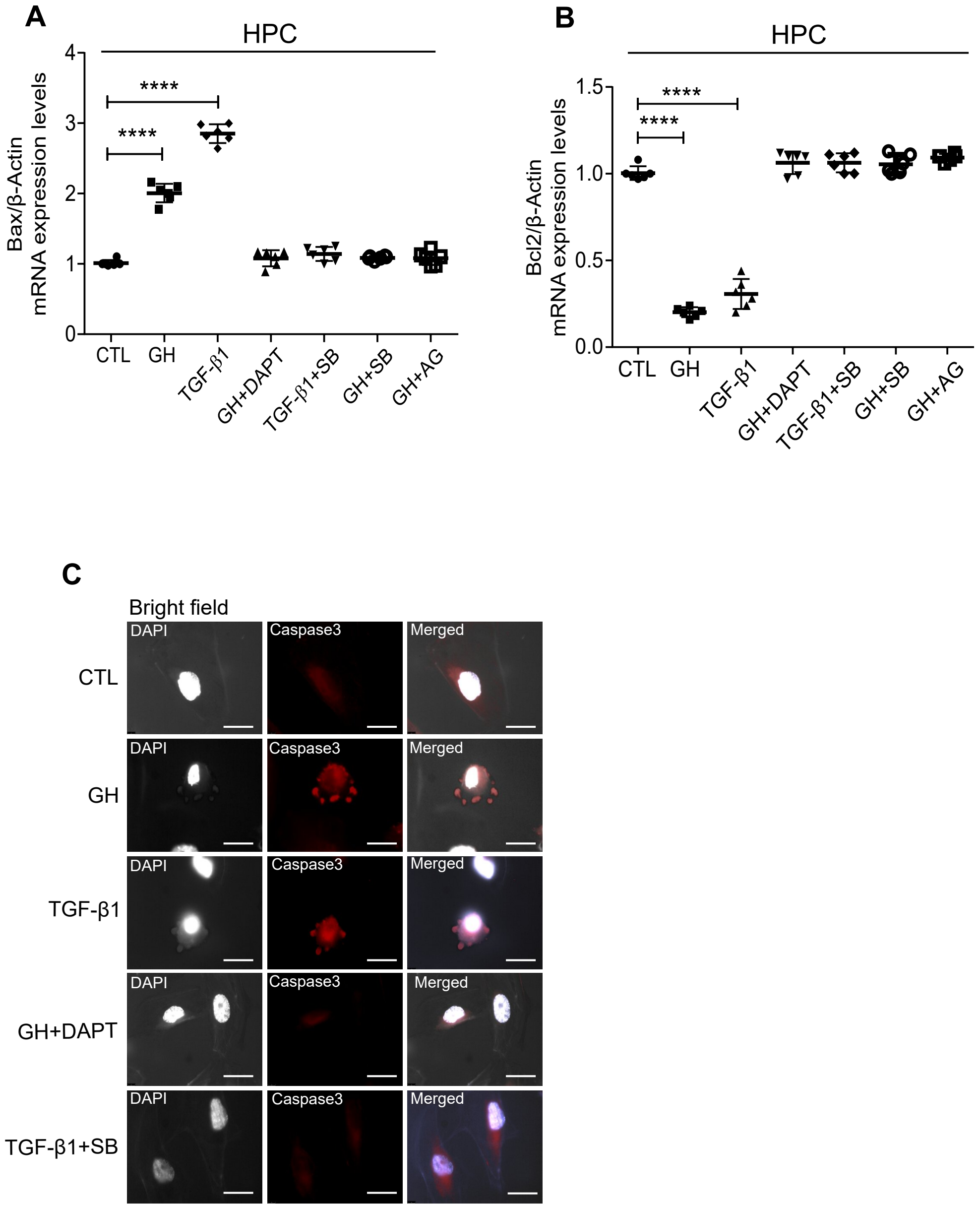

Supplementary Figure 4

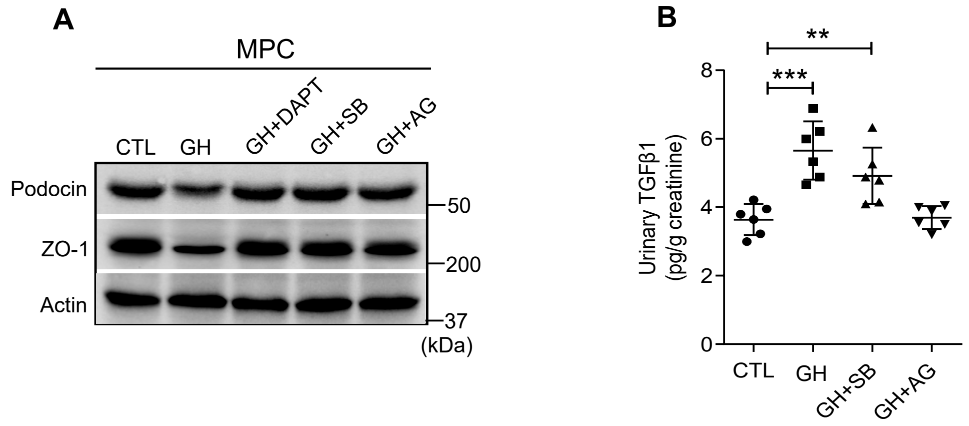
